# Novel Function of Bluetongue Virus NS3 Protein in Regulation of the MAPK/ERK Signaling Pathway

**DOI:** 10.1101/562421

**Authors:** Cindy Kundlacz, Marie Pourcelot, Aurore Fablet, Rayane Amaral Da Silva Moraes, Thibaut Léger, Bastien Morlet, Cyril Viarouge, Corinne Sailleau, Mathilde Turpaud, Axel Gorlier, Emmanuel Breard, Sylvie Lecollinet, Piet A. van Rijn, Stephan Zientara, Damien Vitour, Grégory Caignard

**Author notes:** These authors contributed equally to this work (Damien Vitour, Grégory Caignard). Corresponding authors: Grégory Caignard, UMR VIROLOGIE, Maisons-Alfort, France; Damien Vitour, UMR VIROLOGIE, Maisons-Alfort, France.

## Abstract

Bluetongue virus (BTV) is an arbovirus transmitted by blood-feeding midges to a wide range of wild and domestic ruminants. In this report, we showed that BTV, through its virulence non-structural protein NS3 (BTV-NS3), is able to activate the MAPK/ERK pathway. In response to growth factors, the MAPK/ERK pathway activates cell survival, differentiation, proliferation and protein translation but can also lead to the production of several inflammatory cytokines. By combining immunoprecipitation of BTV-NS3 and mass spectrometry analysis from both BTV-infected and NS3-transfected cells, we identified the serine/threonine-protein kinase B-Raf (BRAF), a crucial player of the MAPK/ERK pathway, as a new cellular interactor of BTV-NS3. BRAF silencing led to a significant decrease of the MAPK/ERK activation by BTV supporting a model where BTV-NS3 interacts with BRAF to activate this signaling cascade. Furthermore, the intrinsic ability of BTV-NS3 to bind BRAF and activate the MAPK/ERK pathway is conserved throughout multiple serotypes/strains but appears to be specific to BTV compared to other members of *Orbivirus* genus. Inhibition of MAPK/ERK pathway with U0126 reduced viral titers, suggesting that BTV manipulates this pathway for its own replication. Therefore, the activation of the MAPK/ERK pathway by BTV-NS3 could benefit to BTV replication by promoting its own viral protein synthesis but could also explain the deleterious inflammation associated with tissue damages as already observed in severe cases of BT disease. Altogether, our data provide molecular mechanisms to explain the role of BTV-NS3 as a virulence factor and determinant of pathogenesis.

**Importance:** Bluetongue Virus (BTV) is responsible of the non-contagious arthropod-borne disease Bluetongue (BT) transmitted to ruminants by blood-feeding midges. Despite the fact that BTV has been extensively studied, we still have little understanding of the molecular determinants of BTV virulence. In this report, we found that the virulence protein NS3 interacts with BRAF, a key component of the MAPK/ERK pathway. In response to growth factors, this pathway promotes cell survival, increases protein translation but also contributes to the production of inflammatory cytokines. We showed that BTV-NS3 enhances the MAPK/ERK pathway and this activation is BRAF-dependent. Our results demonstrate, at the molecular level, how a single virulence factor has evolved to target a cellular function to ensure its viral replication. On the other hand, our findings could also explain the deleterious inflammation associated with tissue damages as already observed in severe cases of BT disease.

## Introduction

Bluetongue Virus (BTV) is the etiologic agent of the non-contagious arthropod-borne disease Bluetongue (BT) transmitted to ruminants by blood-feeding midges of the genus *Culicoides*. It belongs to the *Orbivirus* genus within the *Reoviridae* family, with 27 serotypes currently identified (1) and at least 6 putative new serotypes (2–7). BTV infects a broad spectrum of wild and domestic ruminants even if sheep are the most sensitive species to the disease. During the 20^th^ century BTV was principally circumscribed to tropical and subtropical geographical areas (8). In 2006, BTV serotype 8 (BTV-8, strain 2006) emerged in Northern Europe (9) from which it rapidly spread to Central and Western Europe, causing significant economic losses (mortality, morbidity, reduced production and restrictions in trade of ruminants). Despite the fact that a high vaccination coverage has been achieved across many European countries allowing the control of the BT disease, BTV outbreaks are still a major concern for the World Organisation for Animal Health (OIE), particularly in Europe (1). Clinical signs include hemorrhagic fever, ulcer in the oral cavity and upper gastrointestinal tract, necrosis of the skeletal and cardiac muscle and oedema of the lungs (10). These variabilities in its host range and clinical manifestations are due to several factors related both to the infected hosts and the viral serotypes and strains.

The BTV genome is composed of 10 double-stranded RNA (dsRNA) segments encoding seven structural (VP1 to VP7) and five, or possibly six, non-structural (NS1 to NS4, NS3A and possibly NS5) proteins (11–13). The BT virion is an icosahedral particle organized as a triple-layered capsid. Viral genomic segments are associated with replication complexes containing VP1 (RNA-dependent RNA polymerase), VP4 (capping enzyme including methyltransferase), VP6 (RNA-dependent ATPase and helicase) and enclosed by VP3 (subcore) and VP7 (core) (14). Cell attachment and viral entry involve the two structural proteins of the outer capsid VP5 and the most variable of BTV protein VP2 representing the main target of neutralizing antibodies and determines the serotype specificity (15, 16). Non-structural proteins contribute to the control of BTV replication (17), viral protein synthesis (18), maturation and export from the infected cells (19–23). Initially described for NS4, NS3 has also been shown to counteract the innate immune response, and in particular the type I interferon (IFN-α/β) pathway (13, 24, 25).

NS3 is encoded by the segment 10 and expressed as two isoforms, NS3 and NS3A, the latter being translated from an second in-frame start codon by which the first N-terminal 13 amino acids residues are lacking (26). NS3 proteins are glycoproteins that promote viral release either through its viroporin activity (20) or by budding. This latter implies interactions between NS3 and outer capsid VP2/VP5 proteins (27), and cellular proteins involved in the pathway of endosomal sorting complexes required for transport (ESCRT) (TSG101 and NEDD4-like ubiquitin ligase) and the calpactin light chain p11 (23, 28, 29). Altogether, these reports provide molecular basis of the multifunctional role of BTV-NS3 as a virulence factor and determinant of pathogenesis as also illustrated by other *in vivo* studies using BTV monoreassortants and NS3/NS3A knockout mutants (30–32).

Many viruses can modulate and hijack signaling pathways related to the mitogen-activated protein kinase/extracellular signal-regulated kinase (MAPK/ERK) pathway for more efficient replication (33). In response to extracellular stimuli such as growth factors, several downstream components of the MAPK/ERK pathway including RAS, RAF, MEK1/2 and ERK1/2 are successively activated. Then, ERK1/2 directly or indirectly regulates transcription factors (e.g. Elk1) involved in cell proliferation, differentiation and survival (34, 35) but also cellular factors that control mRNA translation like eukaryotic initiation factor 4E (eIF4E) (36). In 2010, Mortola and colleagues were the first to show the modulation, as activation, of the MAPK/ERK pathway by BTV (37). In contrast to this finding, other studies demonstrated that the phosphorylation of ERK1/2 was reduced (38) or unchanged (39) after BTV infection. The discrepancy of these studies may be due to different virus infection kinetics or differences in viral serotype/strain and/or cell line used. In addition, the molecular mechanisms underlying the potential modulation of this pathway and its possible contribution to the BTV pathogenesis remain to be established. Altogether, these data led us to investigate the functional impact of BTV on the MAPK/ERK signaling pathway.

## Results

### BTV-NS3 activates the MAPK/ERK signaling pathway

To address the question of BTV modulation with the MAPK/ERK pathway, we used a trans-reporter gene assay that measures Elk1 activation by ERK1/2. In this system, Elk1 transcription factor is fused to the DNA binding domain of Gal4 (Gal4-DB) and leads to the expression of the firefly luciferase reporter gene downstream of a promoter sequence containing a Gal4 binding site. Upon stimulation with a growth factor like EGF, Elk1 was activated as assessed by a 5-fold increase of luciferase activity compared to unstimulated HEK-293T cells (Figure 1A; mock control). Cells infected with BTV at different MOIs showed a very strong enhancement in this cellular pathway, even in absence of EGF stimulation (Figure 1A), and in a MOI-dependent manner. Moreover, the inability of a UV-inactivated BTV to activate the luciferase reporter gene indicated that the induction of MAPK/ERK was dependent on viral replication or *de novo* viral protein expression. To determine whether viral protein(s) could be involved in the MAPK/ERK activation, we tested separately all the BTV proteins in our reporter assay. As shown in Figure 1B, only BTV-NS3 is able to strongly activate the MAPK/ERK pathway both in presence or absence of EGF stimulation. Thus, the activation of the MAPK/ERK pathway by BTV notably involves its NS3 viral protein.

**Figure 1.**
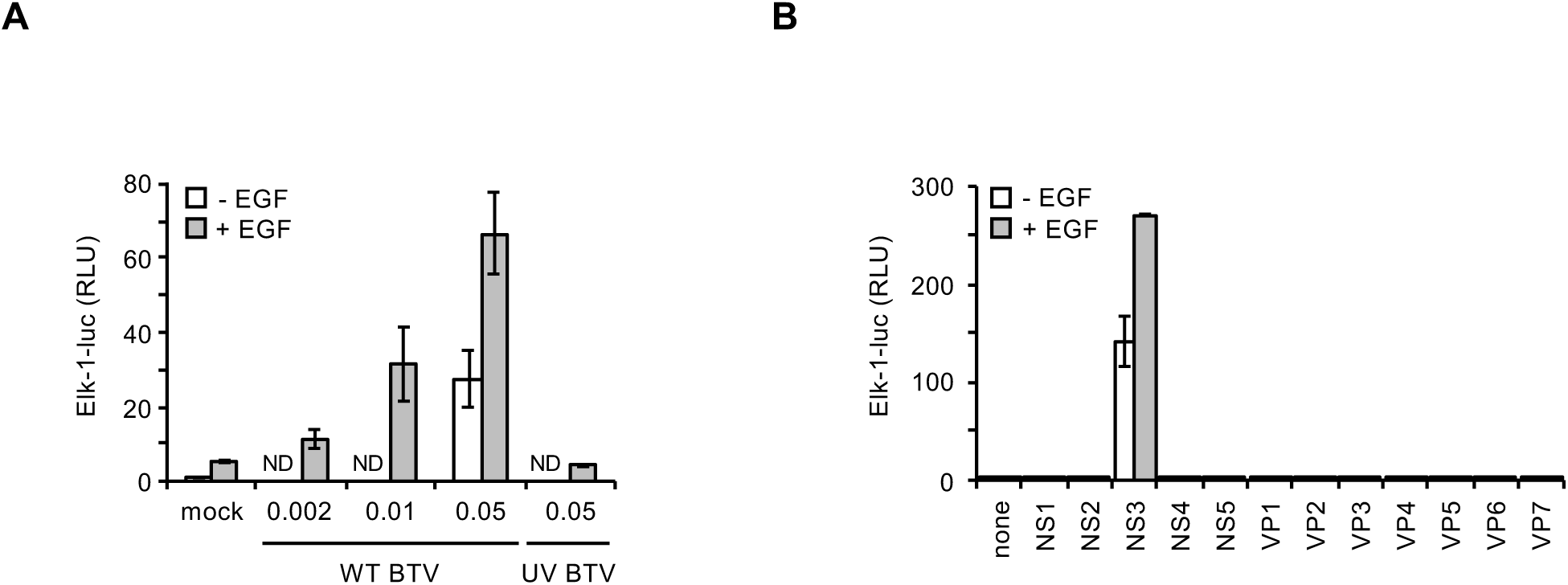
Activation of the MAPK/ERK pathway by BTV-NS3. HEK-293T cells were transfected with pFA2-Elk1 to express Elk1 transcription factor fused to the DNA binding domain of Gal4, pGal4-UAS-Luc that contains the firefly luciferase reporter gene downstream of a promoter sequence containing Gal4 binding site, and pRL-CMV that drives *Renilla* luciferase expression constitutively. In addition to these three plasmids, cells were co-transfected with expression vectors encoding BTV ORFs in fusion with the 3xFLAG tag (B). 12 h after transfection, cells were serum-starved and 6 h later EGF was added at a final concentration of 400 ng/ml (A, B). At the time of EGF stimulation, cells were also infected with WT BTV at indicated MOIs or UV-inactivated BTV (MOI=0.05) (A). After 24 h, relative luciferase activity was determined (A, B). All experiments were achieved in triplicate, and data represent means ± SD. ND: not determined.

### BTV-NS3 interacts with BRAF

As a first approach to understand at molecular level how BTV-NS3 activates the MAPK/ERK pathway, we undertook proteomic analyses after immunoprecipitation of NS3 from either BTV-infected or NS3-transfected HEK-293T cells. The whole purification protocol and LC-MS/MS analyses are presented in the Materials and Methods section. These analyses revealed that BTV-NS3 copurified with BRAF in both BTV-infected and NS3-transfected cells (with Mascot scores of 64 and 104, respectively) whereas BRAF was not detected in the control conditions (mock infected-or empty pCI-neo-3xFLAG transfected-cells). The identified peptides corresponding to BRAF are listed in Table 1.

**Table 1.**
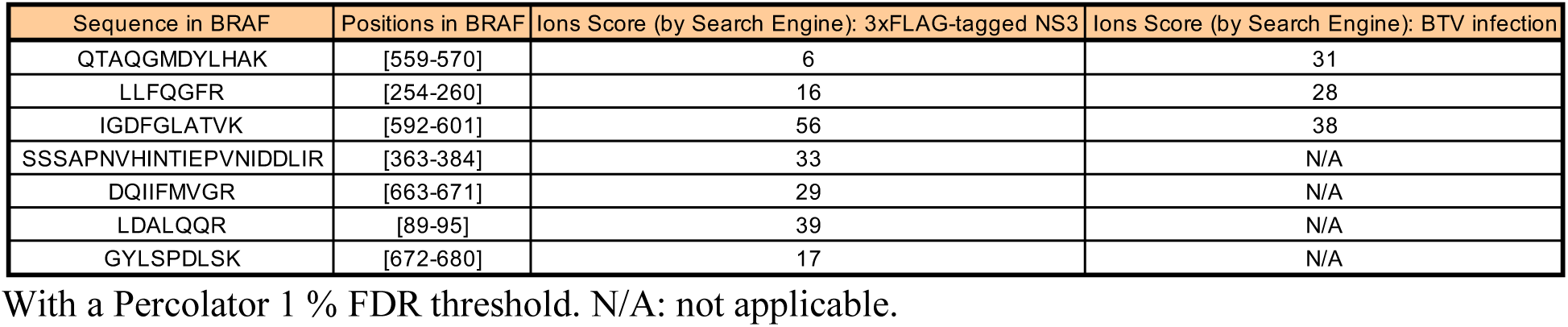
List of the identified peptides corresponding to BRAF.

BRAF, together with ARAF and CRAF, are members of the RAF kinase family that play a central role in regulating the MAPK/ERK signaling pathway. Therefore, binding to BRAF represents a potential molecular mechanism underlying this manipulation that we decided to investigate. To validate this interaction, full-length BTV-NS3 (NS3_FL_) and different fragments of NS3 (Figure 2A) were tested for their ability to interact with endogenous BRAF. To do so, GST-tagged NS3_FL_ or indicated fragments were expressed in HEK-293T cells and purified 48 h later with glutathione-sepharose beads. As shown in Figure 2B, endogenous BRAF copurified only with NS3_FL_.

**Figure 2.**
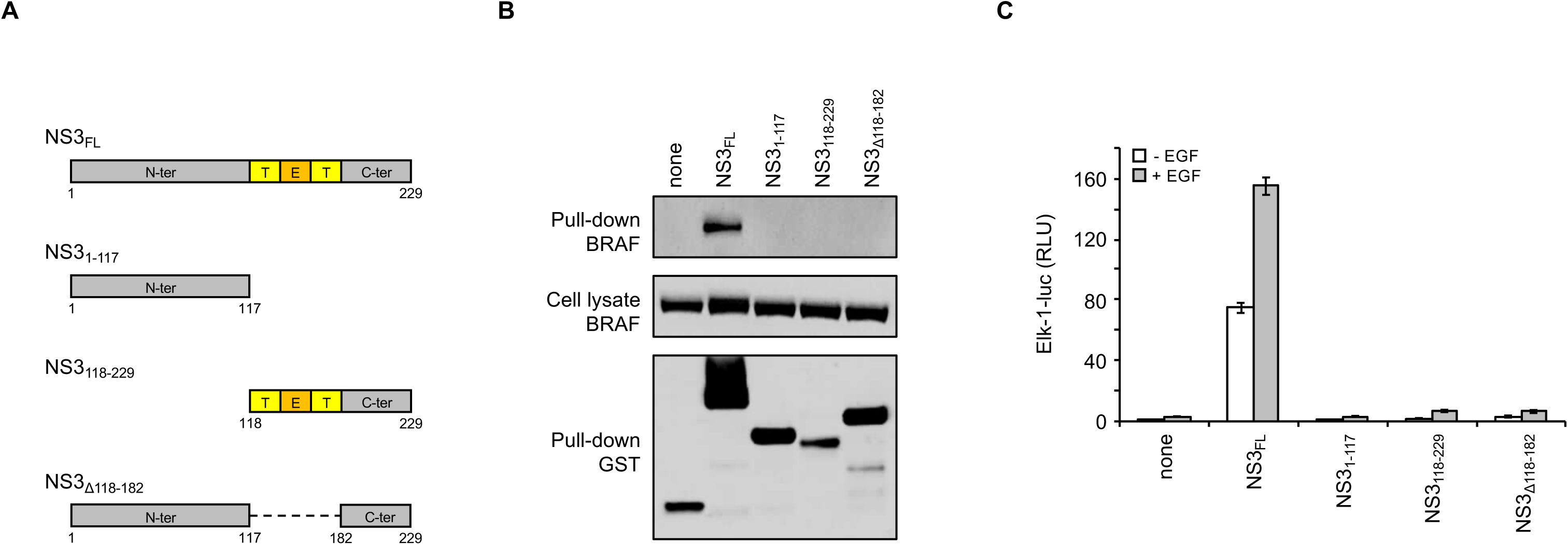
Only full-length BTV-NS3 interacts with BRAF and activates the MAPK/ERK pathway. (A) Schematic representation of BTV-NS3 protein. Deletion fragments of BTV-NS3 were designed. Transmembrane (T) and extracellular domains (E) are indicated. (B) HEK-293T cells were transfected with expression vectors encoding GST alone or fused to the full-length NS3 protein (NS3_FL_) or its indicated fragments, and tested for the interaction with endogenous BRAF. Total cell lysates were prepared 48 h post-transfection (cell lysate; middle panel), and co-purifications of endogenous BRAF were assayed by pull-down using glutathione-sepharose beads (pull-down; upper panel). GST-tagged NS3_FL_ and fragments were detected by immunoblotting using anti-GST antibody (pull-down; lower panel), while endogenous BRAF was detected with a specific antibody. (C) As described in Figure 1, HEK-293T cells were transfected with pFA2-Elk1, pGal4-UAS-Luc and pRL-CMV the expression vector encoding 3xFLAG-tagged NS3_FL_ or fragments as indicated. 12 h after transfection, cells were serum-starved and 6 h later EGF was added at a final concentration of 400 ng/ml. After 24 h, relative luciferase activity was determined. All experiments were achieved in triplicate, and data represent means ± SD. ND: not determined.

Using the same luciferase assay, we showed that only full-length BTV-NS3 is able to enhance Elk1 activation (Figure 2C). In contrast, the indicated fragments of NS3 were unable to do so, consistently with the previous pull-down assays (Figure 2B). Altogether, these results demonstrate that only the full-length BTV-NS3 interacts with BRAF and activates the MAPK/ERK signaling pathway.

### Phosphorylation of ERK1/2 and eIF4E is stimulated by BTV infection and in cells expressing BTV-NS3

To further decipher the impact of BTV-NS3 on the MAPK/ERK signaling pathway, we compared the phosphorylation kinetics of ERK1/2 and eIF4E in HEK-293T cells infected by BTV (Figure 3A) and in cells expressing BTV-NS3 (Figure 3B). HEK-293T cells were infected with BTV (MOI=0.01) and 24 h later, cells were serum-starved for 12 h before being stimulated with EGF. Phosphorylation levels of ERK1/2, determined at 10, 30, 120 min, 6 h and 24 h after stimulation, were markedly and reproducibly higher after BTV infection compared to mock control (Figure 3A). Interestingly, we observed that BTV also induced ERK1/2 phosphorylation even in absence of EGF stimulation. This is reminiscent to what was observed for Elk1 activation (Figure 1A) showing a significant activation of the MAPK/ERK pathway by BTV. In parallel, we also determined the phosphorylation level of the translation initiation factor eIF4E, a downstream target of this pathway that is involved in the control of mRNA translation. In contrast to ERK1/2, the phosphorylation level of eIF4E in BTV infected-cells was increased only at later time points after EGF stimulation whereas p-eIF4E decreased in mock condition (Figure 3A).

**Figure 3.**
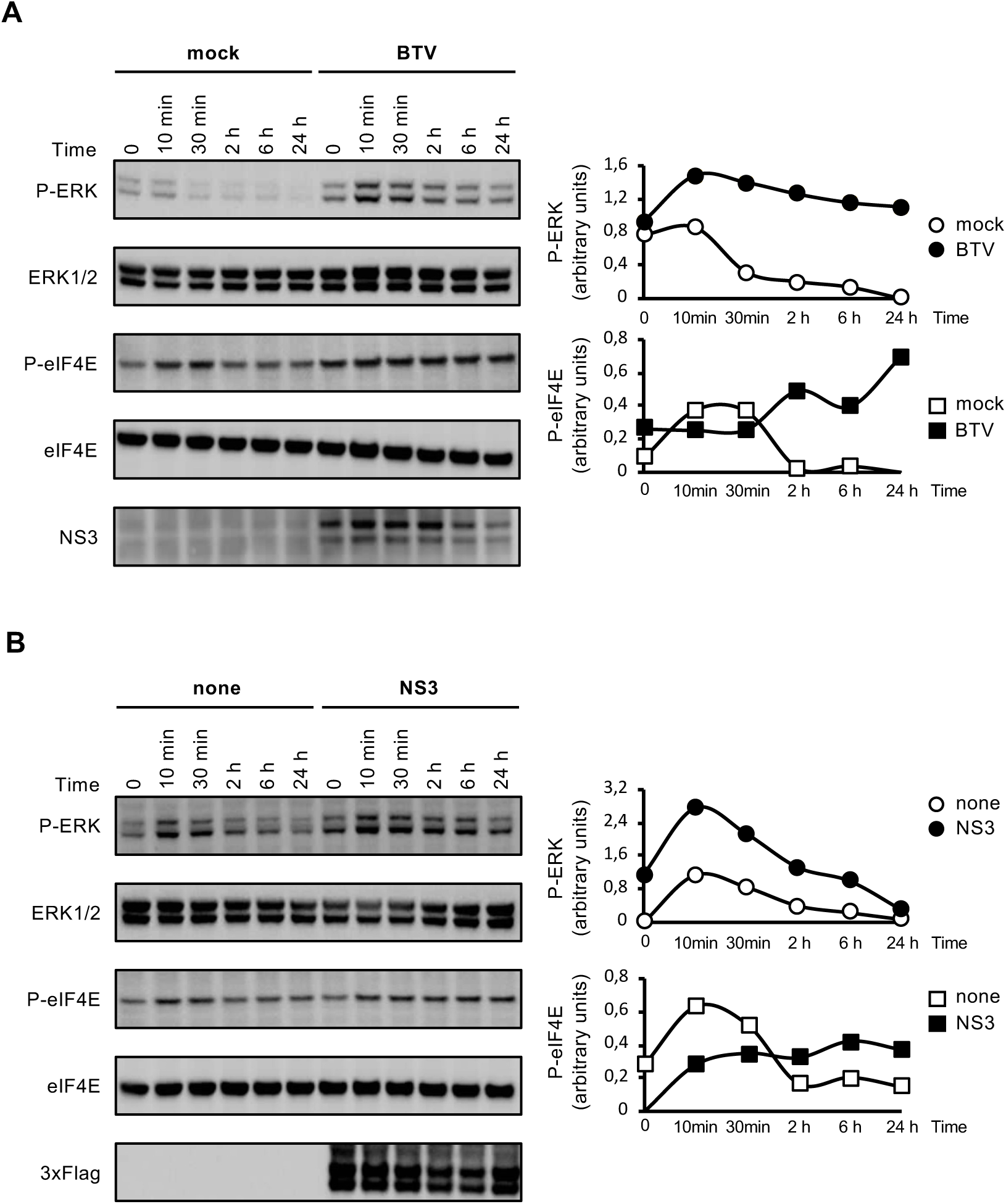
Stimulation of ERK1/2 and eIF4E phosphorylations by BTV infection or BTV-NS3 expression. HEK-293T cells were transfected with an expression vector encoding 3xFLAG-tagged BTV-NS3 or the corresponding empty vector pCI-neo-3xFLAG (B). After 18 h, cells were serum-starved and, when indicated, infected with WT BTV (MOI=0.01) (A). 12 h later, cells were stimulated with 400 ng/ml of EGF. Phosphorylations of ERK1/2 and eIF4E were measured at 10 min, 30 min, 2, 6 and 24 h after EGF stimulation (A, B). BTV infection was confirmed by anti-NS3 immunoblotting (A, lower panel) and expression of 3xFLAG-tagged BTV-NS3 in transfected cells was detected by anti-3xFLAG immunoblotting (B, lower panel). Densitometric analysis of the gels were performed for p-ERK, ERK1/2, peIF4E, eIF4E and the graphs represent the ratio phospho/total. Data presented are representative of at least three independent experiments.

In parallel to the BTV infectious context, cells were transfected with 3xFLAG-tagged BTV-NS3 or a control plasmid. After 24 h, cells were serum-starved and the phosphorylation levels of ERK1/2 and eIF4E were measured at the same time points post-EGF stimulation as before. Like BTV infection, similar phosphorylation kinetics of ERK1/2 and eIF4E were observed for BTV-NS3 expression alone (Figure 3B). In conclusion, these phosphorylation kinetics confirm that either BTV infection or transient expression of BTV-NS3 can both activate the MAPK/ERK pathway.

### BRAF intracellular localization is modified by BTV

Although HEK-293T are highly efficient for transfection and support BTV replication, we aimed to complement our analysis by carrying out similar experiments using a cell line derived from a host naturally infected by BTV. To do so, we measured phosphorylation levels of ERK1/2 in a bovine kidney cell line (MDBK). MDBK cells were serum-starved and infected with BTV at different MOIs for 24 h. As shown in Figure 4A, BTV increased phosphorylation levels of ERK1/2 in a MOI-dependent manner, which is consistent to what was observed for Elk1 activation in HEK-293T cells infected with BTV (Figure 1A).

**Figure 4.**
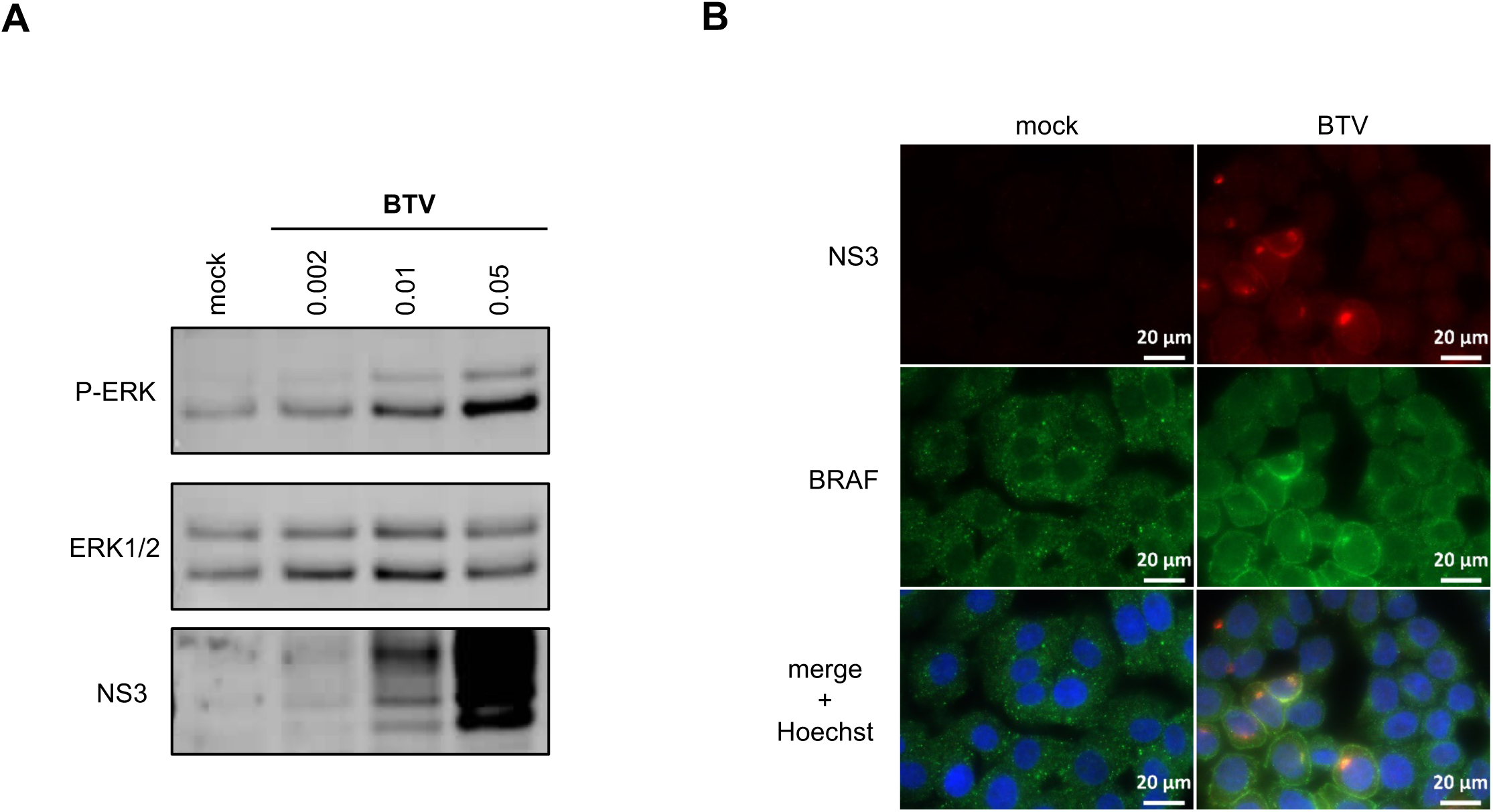
BRAF subcellular localization in the BTV infectious context. (A) MDBK cells were serum-starved and infected with BTV at indicated MOIs for 24 h. Phosphorylated ERK1/2, total ERK1/2 and BTV-NS3 were detected by western blot analysis. (B) MDBK cells were serum-starved and infected with BTV (MOI=0.01). After 24 h, cells were fixed with 4% PFA and labeled with the dye Hoechst 33258 to stain nuclei and with specific antibodies for BRAF and BTV-NS3. Intracellular localization of Hoechst-stained nuclei (blue), endogenous BRAF (green), and BTV-NS3 (red) were visualized by fluorescence microscopy (×63 magnification). Scale bars represent 20 µm.

To determine the potential consequences of NS3-BRAF interaction on their own subcellular localizations, we carried out fluorescence microscopy in MDBK cells. Cells were serum-starved and infected with BTV (MOI=0.01). Then, BTV-NS3 and BRAF localizations were analyzed at 24 h post-infection by fluorescence microscopy (Figure 4B). Firstly, in mock-infected cells, BRAF is present throughout the cytosol with a sparse punctate distribution. As expected, in cells infected by BTV, NS3 is localized in specific cytoplasmic structures evocative of the Golgi apparatus but also at the plasma membrane. Interestingly, we also found that the subcellular distribution of BTV-NS3 matched the relocalization of BRAF in BTV-infected cells. These results demonstrated that BTV-NS3 alters the localization of BRAF, which may contribute to the BTV-activated MAPK/ERK pathway.

### U0126 inhibitor blocks the activation of MAPK/ERK by BTV and alters viral replication

To demonstrate that enhancement of Elk1 activation by BTV-NS3 is completely dependent on ERK1/2 activation, HEK-293T cells were transfected with 3xFLAG-tagged BTV-NS3 or a control plasmid, and 24 h later treated with MEK1/2 inhibitor U0126 (Figure 5A). The U0126 molecule targets MEK1/2 that are directly activated by BRAF proteins (40). As shown in Figure 5A, NS3-induced Elk1 activation was completely inhibited by U0126. To confirm these results, we measured the phosphorylation levels of ERK1/2 and eIF4E in HEK-293T cells treated with U0126 before infection with BTV. As observed for Elk1 activation, U0126 efficiently blocked the phosphorylation of both ERK1/2 and eIF4E after BTV infection (Figure 5B). Interestingly, the presence of U0126 also prevented the expression of BTV-NS3. To test if this inhibitor could have an antiviral effect on BTV replication, HEK-293T cells were treated with MEK1/2 inhibitor U0126 and infected with BTV (MOI=0.01). As shown in Figure 5C, HEK-293T cells treated with U0126 exhibited significant lower viral titers compared to the DMSO control. Altogether, these results suggest that BTV manipulates the MAPK/ERK signaling pathway to increase replication efficiency.

**Figure 5.**
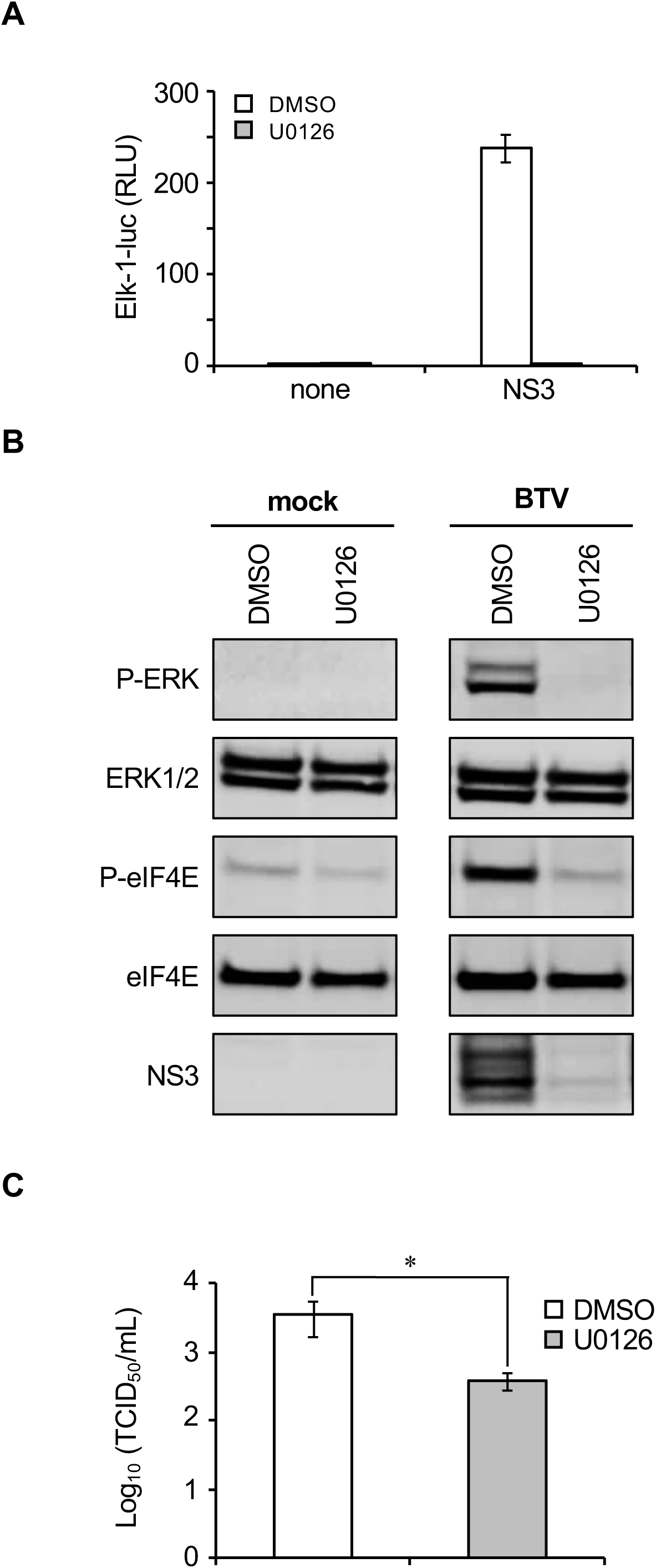
Effects of U0126 inhibitor on BTV-induced MAPK/ERK pathway. (A) As described in Figure 1, HEK-293T cells were transfected with pFA2-Elk1, pGal4-UAS-Luc and pRL-CMV to determine the activation level of MAPK/ERK signaling pathway. In addition to these three plasmids, cells were co-transfected with an expression vector encoding 3xFLAG-tagged BTV-NS3 or the corresponding empty vector pCI-neo-3xFLAG. 12 h after transfection, cells were serum-starved and 6 h later treated with 20 μM of MEK1/2 specific inhibitor U0126 when indicated. After 24 h, relative luciferase activity was determined. All experiments were achieved in triplicate, and data represent means ± SD. (B) HEK-293T cells were serum-starved and infected with BTV (MOI=0.01). At the time of infection, cells were also treated with U0126 inhibitor as indicated. After 24 h, phosphorylations of ERK1/2 and eIF4E were measured and BTV infection was confirmed by anti-NS3 immunoblotting. (C) HEK-293T cells were serum-starved and infected with BTV (MOI=0.01). After 24 h, supernatants were harvested and titrated by TCID50/ml. The experiment was performed in triplicates, and data represent means ± SD. *, p < 0.05.

### BRAF silencing impairs BTV-activated MAPK/ERK pathway

To further confirm the experiments with the MAPK/ERK pharmacologic inhibitor U0126, we used a gene silencing approach targeting BRAF. HEK-293T cells were transfected with BRAF-specific or control non-specific siRNA before infection with BTV (Figure 6A and 6C) or transfection with 3xFLAG-tagged BTV-NS3 (Figure 6B and 6D). In both cases, the reduction of BRAF expression led to a significant decrease of the MAPK/ERK activation as assessed by the luciferase reporter gene assays and anti-pERK/1/2 immunoblotting. These results support a model where BTV-NS3 interaction with BRAF enhances MAPK/ERK pathway activation.

**Figure 6.**
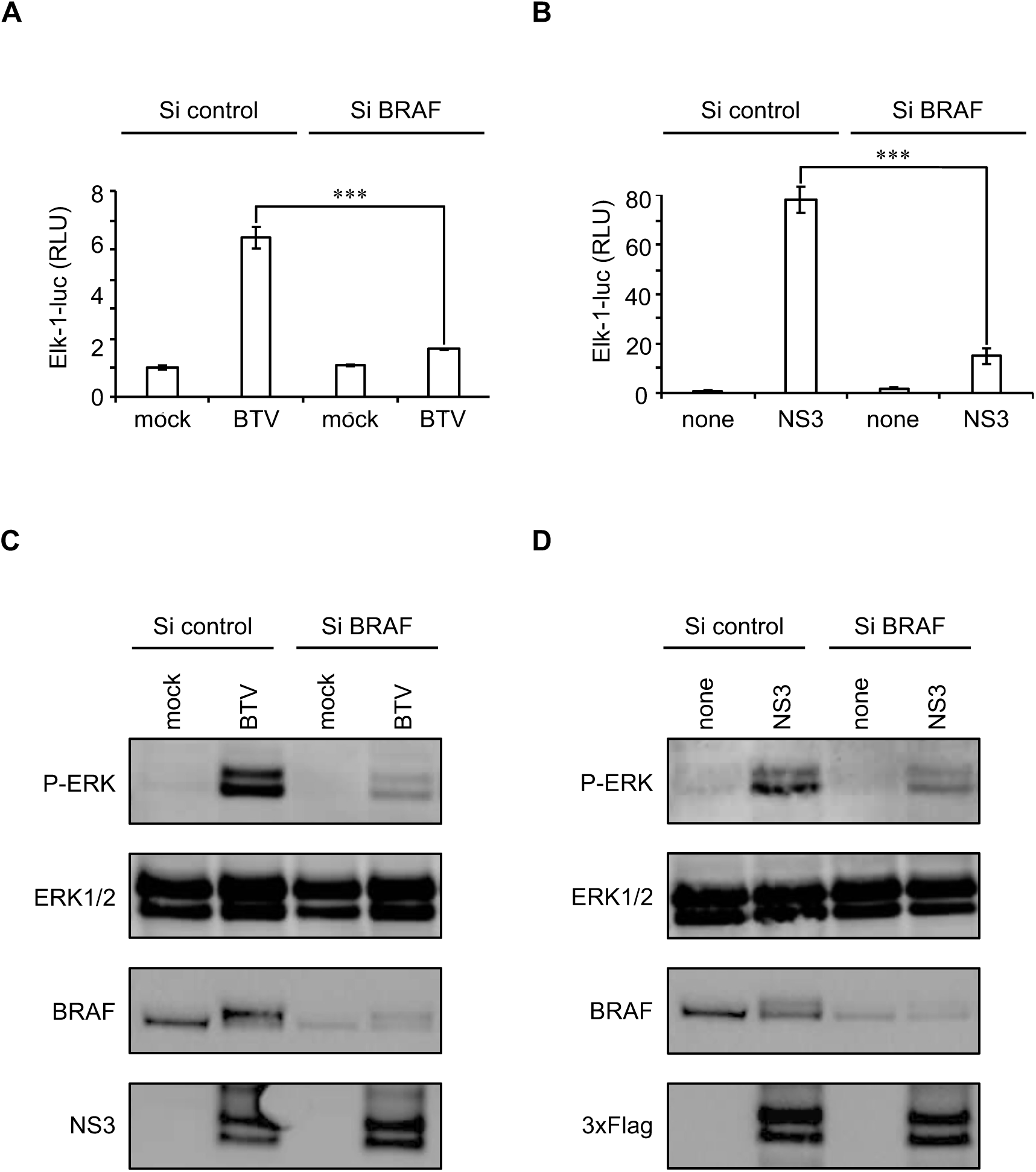
BRAF silencing impairs activation of MAPK/ERK pathway by BTV. (A-D) HEK-293T cells were transfected with non-specific or a BRAF-specific siRNA. One day later, cells were either infected with BTV (MOI=0.01) (A, C) or transfected to express 3xFLAG-tagged BTV-NS3 (B, D) as described in Figure 1. After 24 h, relative luciferase activity was determined (A, B) and cell lysates were analyzed by immunoblotting with antibodies against the indicated proteins (C, D). (A, B) All experiments were achieved in triplicate, and data represent means ± SD. ***, p < 0.0005.

### NS3 interaction with BRAF and activation of the MAPK/ERK signaling pathway are characteristic of BTV

To address the question of the specificity of the BRAF interaction, we first compared NS3 proteins from three serotypes (BTV 1, 8 and 27) in their ability to interact with endogenous BRAF. GST-tagged NS3 from BTV1, 8 and 27 were expressed in HEK-293T cells and purified 48 h later with glutathione-sepharose beads. As shown in Figure 7A, NS3 proteins from BTV1, 8 and 27 have similar binding capacities for BRAF. As a consequence, BTV1-NS3 and BTV27-NS3 were also able to enhance the MAPK/ERK pathway, although the activation by BTV8-NS3 was stronger compared to NS3 proteins from BTV1 and 27 (Figure 7B). Then, NS3 proteins from other members of *Orbivirus* genus, such as epizootic hemorrhagic disease virus (EHDV), African horse sickness virus (AHSV) and equine encephalosis virus (EEV) were also tested for binding to endogenous BRAF. Only BTV-NS3 was able to co-purify with BRAF, demonstrating the specificity of this interaction (Figure 7C). Moreover, only BTV-NS3 highly activates MAPK/ERK pathway, in contrast to EHDV-NS3, AHSV-NS3 and EEV-NS3 (Figure 7D). Thus, the interaction with BRAF and activation of the MAPK/ERK pathway are unique to BTV-NS3.

**Figure 7.**
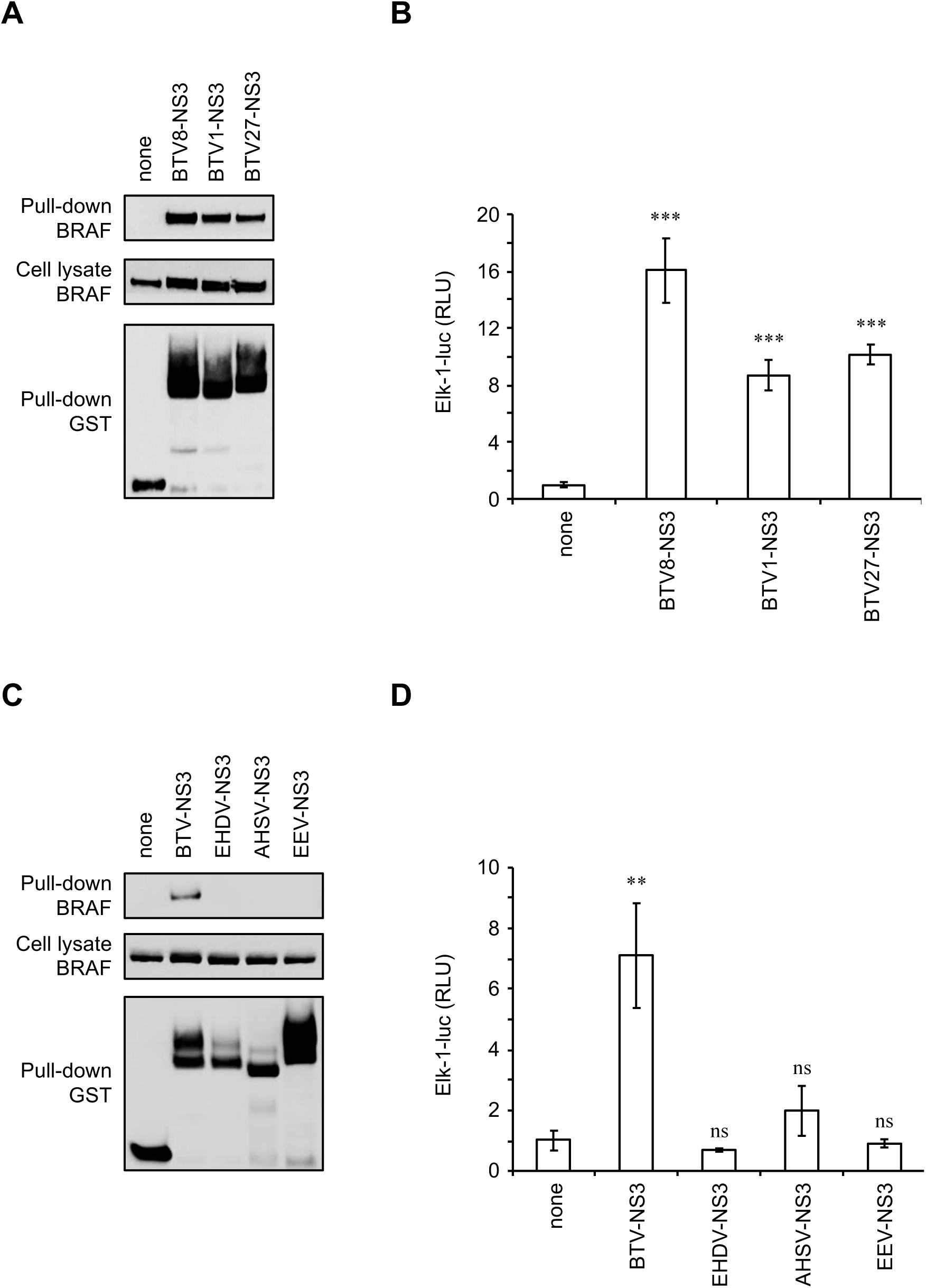
Comparative analysis of MAPK/ERK pathway for NS3 proteins from different orbiviruses. (A, C) HEK-293T cells were transfected with expression vectors encoding GST alone or fused to BTV8-NS3, BTV1-NS3, BTV27-NS3, EHDV-NS3, AHSV-NS3 or EEV-NS3 as indicated, and tested for the interaction with endogenous BRAF. Total cell lysates were prepared 48 h post-transfection (cell lysate; middle panel), and co-purifications of endogenous BRAF were assayed by pull-down using glutathione-sepharose beads (pull-down; upper panel). (B, D) As described in Figure 1, HEK-293T cells were transfected with pFA2-Elk1, pGal4-UAS-Luc and pRL-CMV. In addition to these three plasmids, cells were co-transfected with an expression vector encoding 3xFLAG-tagged BTV8-NS3, BTV1-NS3, BTV27-NS3, EHDV-NS3, AHSV-NS3, EEV-NS3 or the corresponding empty vector pCI-neo-3xFLAG, as indicated. 12 h after transfection, cells were serum-starved for 24 h and relative luciferase activity was then determined. The experiment was performed in triplicate, and data represent means ± SD. *** indicates that differences observed between BTV8-NS3, BTV1-NS3 or BTV27-NS3 and the corresponding control vector pCI-neo-3xFLAG were statistically significant p < 0.0005; ** indicates that differences observed between BTV-NS3 and the corresponding control vector pCI-neo-3xFLAG were statistically significant p < 0.005; ns: non-significant differences between EHDV-NS3, AHSV-NS3 or EEV-NS3 and the corresponding control vector pCI-neo-3xFLAG.

### BTV-NS3 inhibits the induction of IFN-α/β independently of MAPK/ERK signaling

Our team has demonstrated the major role of BTV-NS3 in counteracting the induction of the type I interferon (IFN-α/β) response (24). Interestingly, it has been reported that the activation of the MAPK/ERK pathway could be associated with the inhibition of IFN-α/β synthesis (41). Therefore, we asked if the activation of the MAPK/ERK pathway by BTV-NS3 is required for an efficient control of the IFN-α/β response. Using a luciferase gene reporter assay, we tested NS3_FL_ and its fragments for their capacity to inhibit an IFN-β specific promoter downstream a stimulation with a constitutively active form of RIG-I (NΔRIG-I for N-terminal CARDs of RIG-I). As shown in Figure 8A, only NS3_FL_ fully blocked the IFN-β promoter activity. Moreover, while NS3_118-229_ and NS3_Δ118-182_ fragments were not able to activate Elk1 as previously shown in Figure 2C, both are partially, but significantly, able to block the IFN-β promoter activity (Figure 8A). As a complementary approach, we used the MEK1/2 inhibitor U1026 and measure its impact on the capacity of BTV-NS3 to inhibit the IFN-β promoter activity. As shown in Figure 8B, the U0126 molecule was unable to prevent the antagonist function of BTV-NS3 on the induction of IFN-α/β and thus to rescue a significant activation of the IFN-β promoter. In conclusion, our data demonstrate that the activation of MAPK/ERK by BTV-NS3 does not contribute its antagonist activity on the IFN-α/β synthesis.

**Figure 8.**
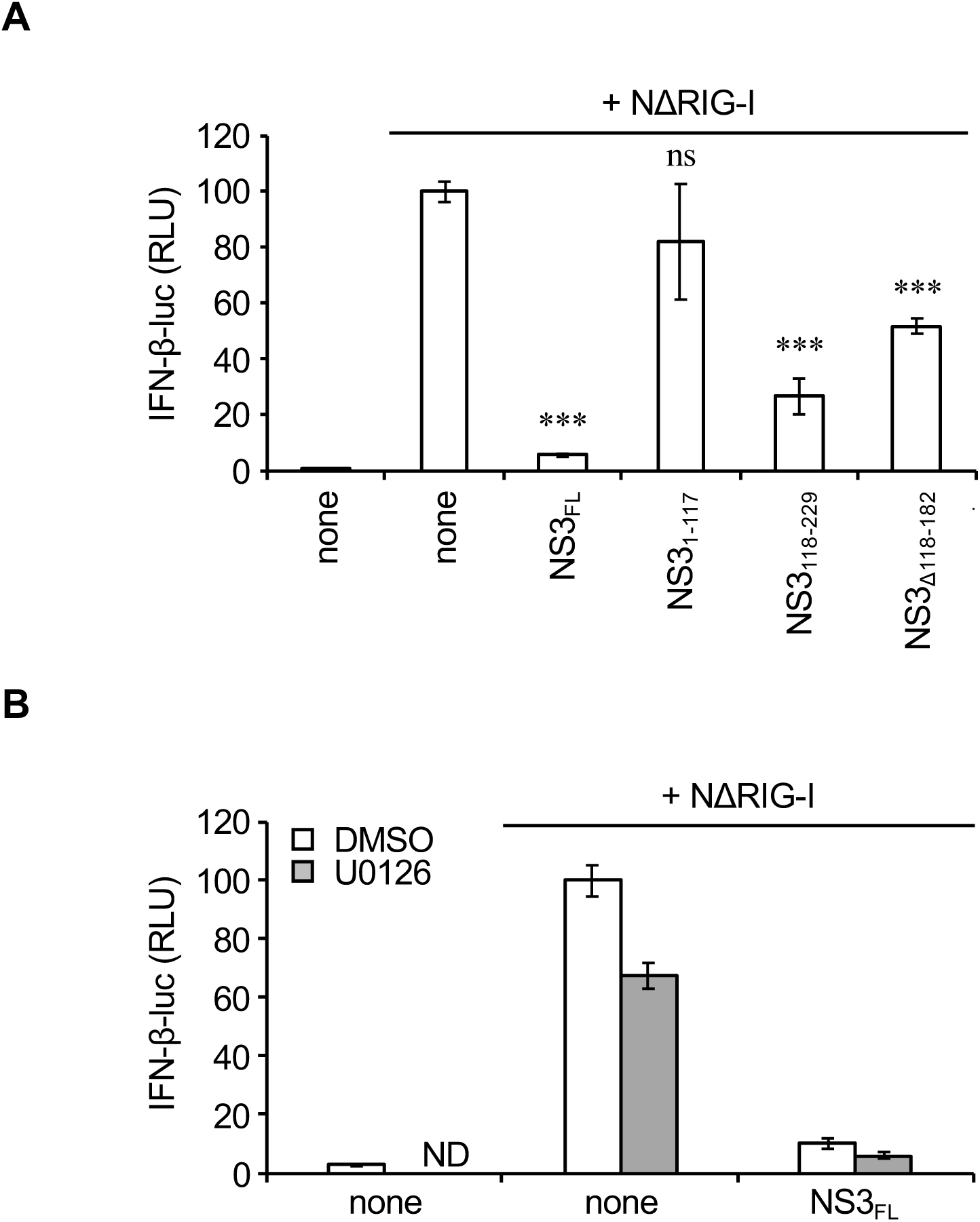
Activation of MAPK/ERK pathway is not related to the inhibition of IFN-α/β signaling by BTV-NS3. (A) HEK-293T cells were co-transfected with IFN-β-pGL3 plasmid that contains the firefly luciferase reporter gene downstream of an IFN-β specific promoter sequence, pRL-CMV reference plasmid, and pCI-neo-3xFLAG expression vectors encoding for 3xFLAG alone or fused to NΔRIG-I (N-terminal CARDs of RIG-I) and BTV-NS3_FL_ or fragments. After 48 h, relative luciferase activity was determined. *** indicates that differences observed between NS3_FL_, NS3_118-229_ or NS3_Δ118-182_ and the corresponding control vector pCI-neo-3xFLAG were statistically significant p < 0.0005; ns: non-significant differences between NS3_1-117_ and the corresponding control vector pCI-neo-3xFLAG. (B) Same experiments as (A) but 24 h after transfection, cells were serum-starved for 6 h and treated with 20 μM of U0126 as indicated. 24 h later, relative luciferase activity was determined. All experiments were performed in triplicates, and data represent means ± SD. ND: not determined.

## Discussion

As obligate intracellular parasites, viruses have evolved multiples strategies to hijack their host cellular machineries and control them to survive, replicate and spread. The MAPK/ERK pathway contains one of the most highly conserved family of serine/threonine kinases from yeast to humans, which regulates a multiplicity of cellular processes including cell survival, proliferation and differentiation, as well as immune and inflammatory responses. Therefore, many viruses have been shown to modulate this pathway for their own benefit (33). Considering that its aberrant activation represents an important step toward carcinogenesis, the modulation of the MAPK/ERK pathway has been initially described for DNA tumor viruses and oncogenic retroviruses. However, non-oncogenic RNA viruses are also able to activate this signaling cascade even if the molecular mechanisms underlying this manipulation, in particular in term of protein-protein interactions, often remain a pending question.

In this report, we show that BTV activates the MAPK/ERK pathway as assessed by Elk1 transactivation and phosphorylation levels of ERK1/2 and eIF4E, which is reminiscent to findings of Mortola and colleagues (37), but we also give the first example of a BTV protein that contributes to this positive regulation. We further demonstrate that both BTV infection and NS3 expression alone also activate the MAPK/ERK pathway in the absence of external stimuli. Moreover, our data provide molecular basis to this activity through the identification of BRAF as a new interactor of BTV-NS3 together with the fact that BRAF silencing impairs BTV-activated MAPK/ERK signaling. Intriguingly, two other studies have shown that BTV does not activate the MAPK/ERK pathway (38, 39). However, their findings are not necessarily in contradiction with our current data. This apparent discrepancy does not appear to depend on factors related to the virus used as we have demonstrated that NS3 proteins from at least three serotypes of BTV have similar abilities to bind BRAF and activate the MAPK/ERK pathway.

In addition, we have confirmed this activation in both human and bovine cells. One possible explanation could be that, in contrast to these previous studies, all of our experiments were performed under starvation conditions, which is likely to be essential since the MAPK/ERK pathway is activated in response to growth factors.

In contrast to BTV, NS3 proteins from EHDV, AHSV and EEV are unable to activate the MAPK/ERK pathway suggesting that BTV-NS3 is likely to be functionally distinct from other *Orbiviruses* NS3 proteins. Although BTV-NS3 shares several domains with EHDV-NS3, AHSV-NS3 and EEV-NS3 (e.g. the amphipathic helix at the N terminus, the late-domain motifs, the extracellular and the two transmembrane domains) (23, 42), these proteins are genetically different. Indeed, the NS3 proteins of AHSV and EEV only share ≈30% of sequence homology with BTV-NS3 whereas the protein sequence is more conserved between BTV-NS3 and EHDV-NS3 (57% of sequence homology), which is consistent with the fact that both BTV and EHDV are transmitted to ruminants. Although we have currently no explanation for this apparent specificity of BTV, these differences in protein sequence could account for their capacity or incapacity to activate the MAPK/ERK pathway. Nevertheless, further investigations will be needed to understand how the MAPK activation could provide an advantage for BTV at molecular/cellular level in comparison to other orbiviruses.

Considering that ERK1/2 regulate more than 160 downstream target factors (43), activation of the MAPK/ERK pathway by BTV could have several consequences on host cell biology. It triggers both NF-κB, c-Jun and STAT1 transcription factors (44, 45) leading to an increased expression of inflammatory factors, in particular cytokines and chemokines that participate to immunity and inflammation such as IL-6 and IL-8 (46–48). Although expression of cytokines and chemokines are critical for developing an efficient antiviral response, their excessive production could also lead to deleterious inflammation by causing tissue damages and, therefore, contributing to disease pathogenesis. Indeed, both *in vivo* and *in vitro* studies have shown that BTV-infected cells secrete numerous pro-inflammatory cytokines, including IL-6 and IL-8 (49–55). A consequence of such production could be an aggravation of the endothelial injury associated with an increased vascular permeability as already observed in severe cases of BT disease (10, 56). Altogether, our data suggest that BTV-NS3 interaction with BRAF enhances MAPK/ERK activation above normal level in infected cells, possibly contributing to a deregulation of the blood vessels permeability to promote BTV replication and spreading.

Besides its effects on immunity and inflammation, activation of the MAPK/ERK pathway has considerable consequences on viral replication as assessed by experiments using MEK1/2 inhibitor U0126 inhibitors. Indeed, it is now well documented that MAPK/ERK pathway inhibition with U0126 highly alters the replication of several viruses such as Influenza virus, Junin virus, Herpes simplex virus 1, Astrovirus, Borna disease virus, Coronavirus, human parainfluenza virus type 3 and Porcine epidemic diarrhea virus (57–64). Similarly, we show in this report that MAPK/ERK pathway inhibition by U0126 prevents BTV protein expression in infected cells resulting in reduced viral titer. It should also be noted that the inhibition of the MAPK/ERK pathway, as assessed by Elk1 transactivation and phosphorylation levels of ERK1/2, was more efficient in cells treated with U0126 than those transfected with the BRAF siRNA (Figure 5A compared to 6B and Figure 5B compared to 6C). These differences could be due to the fact that BRAF expression was not fully blocked following gene silencing (Figures 6C and 6D) and this would explain why NS3 expression, as well as viral titers (data not shown), are not affected in cells transfected with the BRAF siRNA. Our findings also correlate with data obtained with other members of *Reoviridae* family. Indeed, activation of EGF signaling pathway has been correlated with increased reovirus replication and spread through the regulation of multiple steps of the infectious life cycle including viral uncoating and disassembly, viral protein translation and generation of viral progeny (for review, see (65)). Rhesus and group A rotaviruses also activate MAPK/ERK pathway to facilitate viral replication and viral uncoating, respectively (66, 67). A possible link between MAPK/ERK pathway and BTV protein expression could be related to the downstream targets of this pathway which are essential factors of the cellular translational machinery, like eIF4E.

eIF4E is believed to be the least abundant of all initiation factors. Therefore, it can be considered as an excellent target to regulate protein synthesis. Phosphorylation of eIF4E is mediated by MAPK interacting kinase 1 (MNK1), itself mainly activated by ERK1/2 (36). However, MNK1 has also other downstream targets including eukaryotic initiation factor 4G (eIF4G). Once activated, eIF4E interacts with the cap structure and brings translation initiation factors together with the small ribosomal subunit via the scaffold protein eIF4G to initiate cap-dependent mRNA translation (68). Since BTV mRNA are capped like their host counterparts (69, 70), it may be assumed that BTV increases phosphorylation level of eIF4E, thereby stimulating cap-dependent translation, to promote its own viral protein synthesis within infected cells.

Our findings support a model where BTV-NS3 interacts with BRAF to activate MAPK/ERK pathway. To the best of our knowledge, this is the first report demonstrating both BRAF as a target of a viral protein and this interaction as an important step toward a viral manipulation of the MAPK/ERK pathway. We also demonstrate that BTV infection leads to a re-localization of BRAF either at the cell membrane or the Golgi apparatus. Initially described to be mainly regulated at the plasma membrane (71, 72), it has now been clearly established that the control of MAPK/ERK pathway can also occur in the Golgi apparatus (73). Thus, these specific localizations could contribute to the aggregation of NS3-BRAF complexes to enhance MAPK/ERK signaling. Further analyses with confocal microscopy will be needed to confirm the colocalization between NS3 and BRAF and additional biochemical investigations are still required to decipher how this viral protein activates BRAF and the downstream events of this pathway. Altogether, NS3 interactions with BRAF represents a potential target for the development of antiviral molecules against BTV.

## Materials and Methods

### Cell lines and viral infections

HEK-293T and MDBK cells were maintained in Dulbecco's modified Eagle's medium (DMEM; Gibco-Invitrogen) containing 10% fetal bovine serum, penicillin, and streptomycin at 37°C and 5% CO_2_. BTV8 WT strain was amplified and titrated on BSR-T7. Inactivated virus was prepared by exposing live virus to 254-nm UV as previously described (74). Serum-free medium was used as an inoculum for BTV-infected cells. BTV infection was analyzed at the indicated time points using a specific NS3 antibody (kindly provided by Dr. Frederick Arnaud) (75) and visualized by fluorescence microscopy or western blotting.

### Plasmid DNA constructs

ORFs-encoding sequences from BTV8 WT strain (isolated in the French Ardennes in 2006 (76)), BTV1 WT strain (isolated in France in 2008 (77)), BTV27 WT strain (isolated in Corsica in 2014 (78)), EHDV6 WT strain (isolated in Reunion Island in 2009 (79)), AHSV4 WT strain (isolated in Morocco in 1990 (80)) and EEV3 WT strain (isolated in South Africa in 1974 (81)) were amplified by RT-PCR (Roche) from purified infected-cell RNAs. Amplification was performed using ORF specific primers flanked with the following Gateway cloning sites: 5′-ggggacaactttgtacaaaaaagttggc and 5′-ggggacaactttgtacaagaaagttgg. PCR products were cloned by *in vitro* recombination into pDONR207 (Gateway system; Invitrogen). ORF coding sequences were subsequently transferred by *in vitro* recombination from pDONR207 into different Gateway-compatible destination vectors (see below) following manufacturer's recommendation (LR cloning reaction, Invitrogen). In mammalian cells, GST-tag and 3xFLAG-tag fusions were achieved using pDEST27 (Invitrogen) and pCI-neo-3xFLAG vector, respectively (82). An expression vector pNRIG-I carrying genes for the constitutively active N-terminal CARDs of RIG-I (NRIG-I) has been used to stimulate the luciferase reporter gene downstream of an IFN-β specific promoter sequence as previously described (84).

### Luciferase reporter gene assay

HEK-293T were plated in 24-well plates (5×10^5^ cells per well). One day later, cells were transfected with either pFA2-Elk1 (0.3 µg/well; Stratagene) and pGal4-UAS-Luc plasmids (0.3 µg/well; provided by Dr. Yves Jacob) or IFN-β-pGL3 (0.3 µg/well; Stratagene) together with pRL-CMV reference plasmid (0.03 µg/well; Promega). Cells were simultaneously co-transfected with 0.3 µg/well of the empty pCI-neo-3xFLAG expression vector or encoding viral proteins as specified. 12 h after transfection, cells were serum-starved for 6 h then stimulated with EGF (Sigma) at 400 ng/ml. When specified, at the time of EGF stimulation, cells were infected by BTV with the indicated MOI. After 24 h, cells were lysed, and both firefly and *Renilla* luciferase activities in the lysate were determined using the Bright-Glo and Renilla-Glo Luciferase Assay System (Promega), respectively. Reporter activity was calculated as the ratio of firefly luciferase activity to reference *Renilla* luciferase activity. All graphs represent the mean, and include error bars of the standard deviation.

### Statistical analyses

p-values are a result of unpaired two-tailed Student’s T test. Differences were considered to be significant if P value <0.05 (*) or <0.005(**) or <0.0005(***).

### Co-affinity purification experiments

To perform co-affinity purification experiments coupled to mass spectrometry analyses, HEK-293T cells were either infected with BTV8 WT strain or transfected with pCI-neo-3xFLAG expression vectors encoding for 3xFLAG alone or fused to BTV-NS3_FL_. Briefly, 2×10^6^ HEK-293T cells were dispensed in each well of a 6-well plate (3 wells per condition), and 24 h later, infected (MOI=0.1) or transfected with 1 µg of each plasmid DNA (JetPRIME; Polyplus). 18 h post-infection or 24 h post-transfection, cells were washed in PBS, then resuspended in lysis buffer (20 mm MOPS-KOH pH7.4, 120 mm of KCl, 0.5% Igepal, 2 mm β-Mercaptoethanol), supplemented with Complete Protease Inhibitor Cocktail (Roche). Cell lysates were incubated on ice for 20 min, then clarified by centrifugation at 14,000×g for 10 min. Protein extracts were incubated for 3 h on a spinning wheel at 4°C with 40 µl of Protein G Sepharose beads (Roche) and 2.5 µg of the specific BTV-NS3 antibody. Beads were then washed with ice-cold lysis buffer 3 times for 5 minutes on a spinning wheel.

For other co-affinity purification experiments, ORFs encoding NS3 or fragments were transferred from pDONR207 to pDEST27 expression vector (Invitrogen) to achieve GST fusion. HEK-293T cells were transfected with 300 ng of each plasmid DNA per well. Two days post-transfection, cells were collected in PBS and then incubated on ice in lysis buffer for 20 min and clarified by centrifugation at 14,000×g for 10 min. For pull-down analysis, protein extracts were incubated for 2 h at 4°C with 30 µl of glutathione-sepharose beads (Amersham Biosciences) to purify GST-tagged proteins. Beads were then washed with ice-cold lysis buffer 3 times for 5 minutes and proteins were recovered by boiling in denaturing loading buffer (Invitrogen).

### LC-MS/MS analyses

Co-immunoprecipitation beads from two independent biological replicates were eluted in Laemmli buffer and run on a 4-12 % acrylamide gel (Invitrogen) and proteins were stained with Coomassie blue (Biorad). The experiment was reproduced one time to obtain two independent biological replicates of the total experiment. For both experiments, three gel plugs were cut for each condition. Plugs were reduced with 10 mM DTT, alkylated with 55 mM iodoacetamide (IAA) and incubated with 20 µl of 25 mM NH_4_HCO_3_ containing 12.5 µg/ml sequencing-grade trypsin (Promega, France) overnight at 37°C. The resulting peptides were sequentially extracted from the gel with 30 % acetonitrile, 0.1 % formic acid and 70 % acetonitrile. Digests were pooled (three per condition) according to the different experimental conditions. Peptides mixtures were analyzed by a Q-Exactive Plus coupled to a Nano-LC Proxeon 1000 (both from Thermo Scientific). Peptides were separated by chromatography with the following parameters: Acclaim PepMap100 C18 pre-column (2 cm, 75 µm i.d., 3 µm, 100 Å), Pepmap-RSLC Proxeon C18 column (50 cm, 75 µm i.d., 2 µm, 100 Å), 300 nl/min flow rate, a 98 min gradient from 95 % solvent A (water, 0.1 % formic acid) to 35 % solvent B (100 % acetonitrile, 0.1% formic acid). Peptides were analyzed in the Orbitrap cell, at a resolution of 70,000, with a mass range of m/z 375-1500. Fragments were obtained by higher-energy collisional dissociation (HCD) activation with a collisional energy of 28 %. MS/MS data were acquired in the Orbitrap cell in a Top20 mode, at a resolution of 17,500. For the identification step, all MS and MS/MS data were processed with the Proteome Discoverer software (Thermo Scientific, version 2.2) and with the Mascot search engine (Matrix Science, version 5.1). The mass tolerance was set to 6 ppm for precursor ions and 0.02 Da for fragments. The following modifications were allowed: oxidation (M), phosphorylation (ST), acetylation (N-term of protein), carbamidomethylation, (C). The SwissProt database (02/17) with the *Homo sapiens* taxonomy and a database including all the viral proteins encoded by BTV were used in parallel. Peptide identifications were validated using a 1% FDR (False Discovery Rate) threshold calculated with the Percolator algorithm. Proteins with at least 2 unique peptides were considered. Identified proteins were considered as potential partners of BTV-NS3 if no identifications were reported in the control condition (mock infected-or empty pCI-neo-3xFLAG transfected-cells).

### Western blot analysis

Purified complexes and protein extracts were resolved by SDS-polyacrylamide gel electrophoresis (SDS-PAGE) on 4–12% NuPAGE Bis–Tris gels with MOPS running buffer and transferred to a nitrocellulose membrane (Invitrogen). Proteins were detected using standard immunoblotting techniques. 3×FLAG- and GST-tagged proteins were detected with a mouse monoclonal HRP-conjugated anti-3×FLAG antibody (M2; Sigma-Aldrich) and a rabbit polyclonal anti-GST antibody (Sigma-Aldrich), respectively. Specific antibodies (all from Cell Signaling) were used to detect endogenous BRAF (clone D9T6S), phospho-ERK1/2 (clone-E10), ERK1/2, phospho-eIF4E (Ser209), eIF4E. Secondary anti-mouse and anti-rabbit HRP-conjugated antibodies were purchased from Invitrogen. Densitometric analysis of the gels was performed using the ImageJ program.

### Immunofluorescence assays

24-well plates (ibidi µ-plates, BioValley) were seeded with MDBK cells (1×10^5^ cells per well). One day later, cells were serum-starved and then infected with BTV (MOI=0.01). 18 h post-infection, cells were washed 3 times with PBS and then incubated with a 4% PFA solution (Electron microscopy sciences) for 30 min at room temperature (RT), and then treated with PBS-Glycine (0.1 M) and PBS-Triton (0.5%) for 5 min/each at RT to quench and permeabilize the cells, respectively. Cells were washed 3 times with PBS and incubate for 1 h with a PBS-BSA 1% blocking solution. Finally, cells were incubated with specific BRAF (Sc-5284; Santa Cruz) and NS3 antibodies for 2 h at RT and then incubated for 1 h at RT in a PBS-BSA 1% solution containing the dye Hoechst 33258 and secondary antibodies (anti-rabbit/A11035 and anti-mouse/A11029; Thermofisher). Preparations were visualized using an Axio observer Z1 fluorescence inverted microscope (Zeiss). Each experiment was repeated at least 3 times.

### MAPK inhibitors and BRAF silencing

When specified, cells were treated with MEK1/2 specific inhibitor U0126 (20 μM final; Promega). Transfections with siRNAs were performed using JetPRIME (Polyplus) according to the manufacturer’s instructions. Control non-specific and BRAF specific siRNAs (SMARTpool: ON-TARGETplus Human BRAF SiRNA) were purchased from Dharmacon and were used at a final concentration of 50 nM.

## Acknowledgements

This work was supported by the Laboratoire d'Excellence “Integrative Biology of Emerging Infectious Diseases” (grant no. ANR-10-LABX-62-IBEID). The LC-MS/MS equipment was funded by the Region Ile-de-France (SESAME), the Paris-Diderot University (ARS) and the CNRS. The funders had no role in study design, data collection and interpretation, or the decision to submit the work for publication.

We would like to thank Dr. Frederick Arnaud, Dr. Marc Therrien, Dr. Pierre-Olivier Vidalain, Dr. Maxime Ratinier, Dr. Virginie Doceul and all members of the UMR 1161 Virology for fruitful discussions. We thank Dr. Yves Jacob and Dr. Frederick Arnaud for providing the pGal4-UAS-Luc plasmid and the NS3 antibody, respectively.

